# SonHi-C: a set of non-procedural approaches for predicting 3D genome organization from Hi-C data

**DOI:** 10.1101/392407

**Authors:** Kimberly MacKay, Mats Carlsson, Anthony Kusalik

## Abstract

1

**Background:** Many computational methods have been developed that leverage the results from biological experiments (such as Hi-C) to infer the 3D organization of the genome. Formally, this is referred to as the 3D genome reconstruction problem (3D-GRP). None of the existing methods for solving the 3D-GRP have utilized a non-procedural programming approach (such as constraint programming or integer programming) despite the established advantages and successful applications of such approaches for predicting the 3D structure of other biomolecules. Our objective was to develop a set of mathematical models and corresponding non-procedural implementations for solving the 3D-GRP to realize the same advantages.

**Results:** We present a set of non-procedural approaches for predicting 3D genome organization from Hi-C data (collectively referred to as SonHi-C and pronounced “sonic”). Specifically, this set is comprised of three mathematical models based on constraint programming (**CP**), graph matching (**GM**) and integer programming (**IP**). All of the mathematical models were implemented using non-procedural languages and tested with Hi-C data from *Schizosaccharomyces pombe* (fission yeast). The **CP** implementation could not optimally solve the problem posed by the fission yeast data after several days of execution time. The GM and IP implementations were able to predict a 3D model of the fission yeast genome in 1.088 and 294.44 seconds, respectively. These 3D models were then biologically validated through literature search which verified that the predictions were able to recapitulate key documented features of the yeast genome.

**Conclusions:** Overall, the mathematical models and programs developed here demonstrate the power of non-procedural programming and graph theoretic techniques for quickly and accurately modelling the 3D genome from Hi-C data. Additionally, they highlight the practical differences observed when differing non-procedural approaches are utilized to solve the 3D-GRP.

## 2 Background

Within the nucleus, a cell’s genetic information undergoes extensive folding and reorganization throughout normal physiological processes. Just like in origami where the same piece of paper folded in different ways allows the paper to take on different forms and potential functions, it is possible that different genomic organizations are related to various nuclear functions. Until recently, it has been extremely difficult to comprehensively investigate this relationship due to the lack of high-resolution and high-throughput techniques for identifying genomic organizations. The development of a technique called Hi-C (based on chromosome conformation capture) [36] has made it possible to detect the complete set of genomic regions in close physical proximity. This proximity is often referred to as an “interaction” between two genomic regions. These interactions can be categorized as either intra-chromosomal (*cis*) interactions or inter-chromosomal (*trans*) interactions (Figure 1).

**Figure 1:**
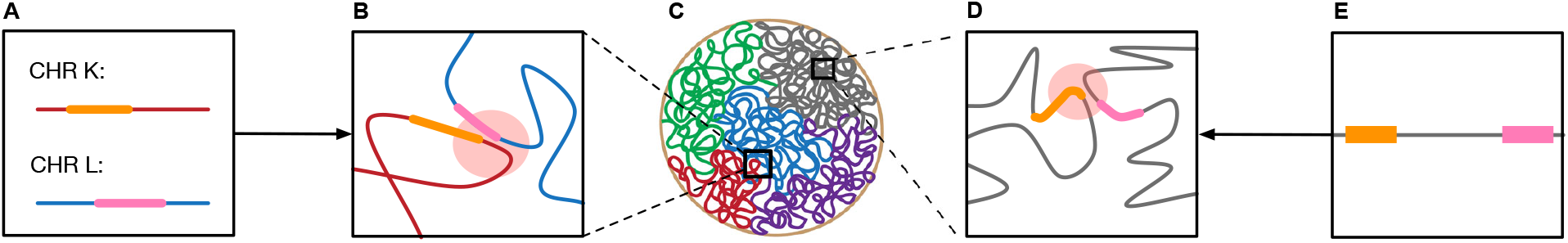
A representation of the DNA-DNA interactions that can occur within the 3D genome structure. Panels give the following representations. A: the linear locations of the genes undergoing a *trans*-interaction between two hypothetical chromosomes, K and L. B: a *trans*-interaction. C: a nucleus with the coloured lines representing the separate chromosomes from Babaei et al. [3]. D: a *cis*-interaction. These genes might be linearly “distant” but still have a detectable interaction in 3D space. E: the linear locations of the genes that are undergoing a 3D *cis*-interaction. The orange and pink regions in panels A, B, D and E are examples of possible gene locations. The red circles in panels B and D represent the genomic regions involved in an interaction.

It is currently unknown whether the 3D genomic organization drives various nuclear functions or *vice versa*, Alterations in the 3D organization of chromosome territories have been demonstrated in a wide variety of cellular processes, including differentiation [31], serum response [39], therapeutic response [21, 40] and response to DNA damage [41]. The unique spatial organization of the genome that is seen under these different cellular conditions is hypothesized to be a crucial mechanism driving various nuclear and cellular functions. It has been theorized that this dynamic organization of the genome may be driven by the global regulation of gene expression [1, 7, 11]. Therefore, the identification of distinct genome interactions may highlight novel mechanisms responsible for organism health and development.

Hi-C [36] is a biological technique that utilizes next generation sequencing technologies to detect regions of the genome that are interacting in 3D space. These regions may be located on different chromosomes or distally on the same chromosome. An overview of the experimental procedure is depicted in Figure 2. Briefly, (1) cells are fixed with formaldehyde in order to covalently cross-link genomic regions that are in close 3D proximity. (2) The crosslinked fragments are then digested with a restriction enzyme to remove the potentially large non-interacting interconnecting segments of DNA. (3) The sticky ends generated through the restriction digest in step (2) are filled in with biotinylated nucleotides. (4) Digested fragments are ligated together. (5) The initial cross-linking is removed, resulting in DNA fragments that represent the two genomic regions that form an interaction. (6) The biotinylated products are purified using streptavidin beads allowing for the detection of fragments that were cut by restriction enzymes. (7) Paired-end sequencing is then performed and the resultant reads are mapped to a reference genome using a Hi-C specific read mapper [2].

**Figure 2:**
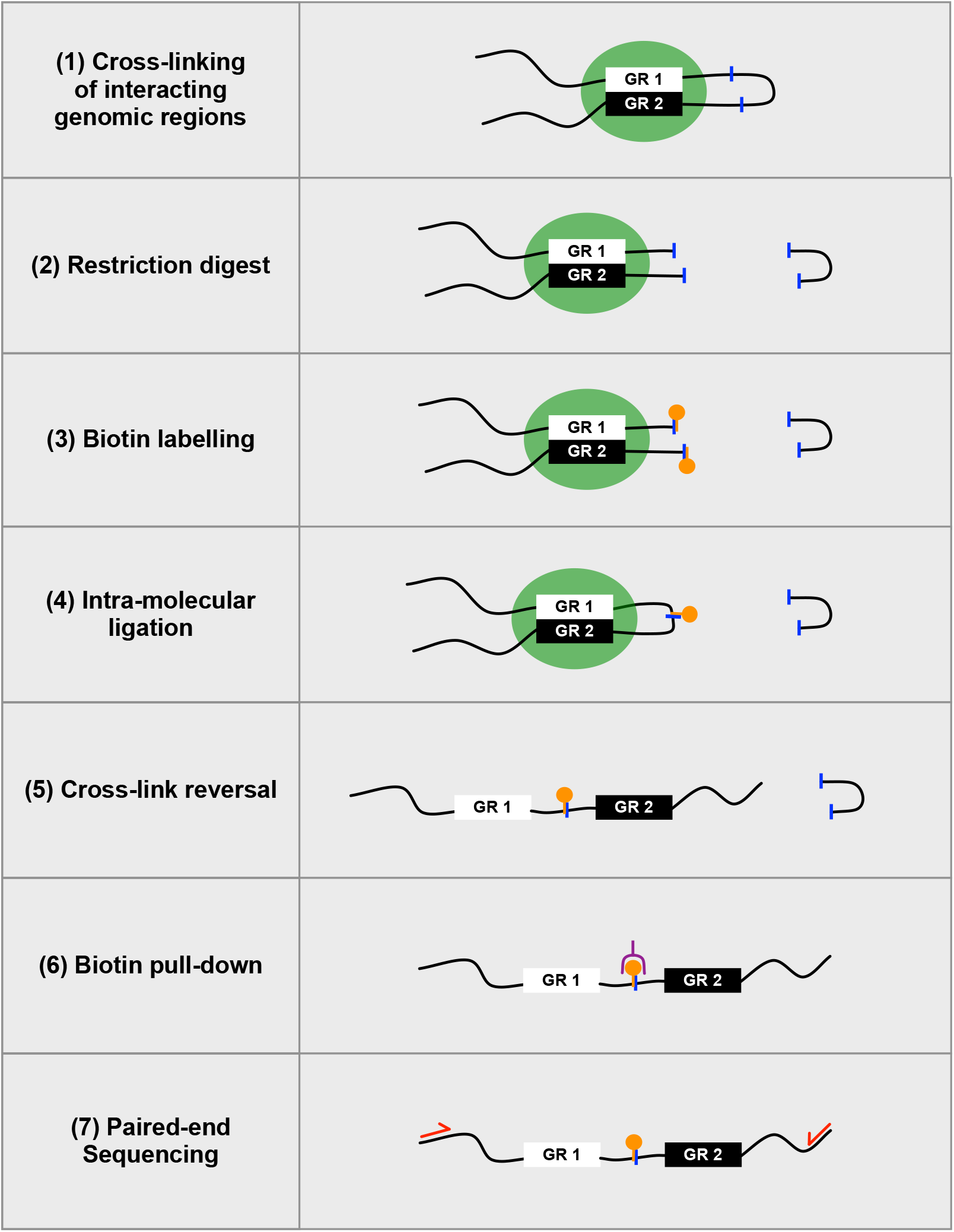
A simplified overview of the Hi-C protocol adapted from reference [36]. GR stands for “genomic region”. The blue lines represent the location of a restriction enzyme cut site; green circles, a pair of genomic regions being chemically cross-linked together; orange circles, biotin; and red arrows, the primers that are required for paired-end sequencing. The purple symbol represents a streptavidin bead that can be used to purify molecules with a biotin label.

Mapping the raw data of a Hi-C experiment to a reference genome results in the generation of a *N × N* matrix (a whole-genome contact map) where N is the number of “bins” which represent linear regions of genomic DNA. In general, the number of genomic bins is approximately equal to the total genome size divided by the Hi-C experimental resolution. Whole-genome contact maps are characteristically sparse and symmetric along the diagonal. Each cell (*A_i, j_*) of a hypothetical whole-genome contact map (**A**) records the count of how many times the genomic bin *i* was found to interact with the genomic bin *j*. These counts are often referred to as the frequency of the interaction between *A_i_* and *A_j_* (or interaction frequency). Inherent systematic biases within the whole-genome contact map are dampened by normalizing the interaction frequencies. Typically, an iterative correction and eigenvector decomposition (ICE) [24] or Knight-Ruiz (KR) [27, 35] normalization are/is applied to the raw data resulting in fractional interaction frequencies.

These normalized whole-genome contact maps can be used to infer the 3D organization of the genome. The process of predicting a model of the 3D genomic organization from a contact map is known as the 3D genome reconstruction problem (3D-GRP) [50]. Typically this is done by converting the normalized interaction frequencies into a set of corresponding pairwise Euclidean distances. In general it is assumed that a pair of genomic regions with a higher interaction frequency will often be closer in 3D space than a pair of genomic regions with a lower interaction frequency [14, 23, 34]. Most computational tools for solving the 3D-GRP then take the predicted pairwise Euclidean distances and produce a visualization of the 3D genome by modelling the chromatin fibre as a polymer [52]. In general, most existing programs can be broadly classified as either (1) consensus or (2) ensemble methods. Consensus methods generate a single population-averaged genomic model that best represents the whole-genome contact map, while ensemble methods produce a collection of genome models that represent the inherent heterogeneity of genome organizations within a population of cells [33].

None of the current methods to solve the 3D-GRP have used a non-procedural approach (such as constraint programming (CP), integer programming (IP) or mixed-integer programming (MIP)) even though non-procedural approaches have been successfully used to predict the 3D structure of other biomolecules [5, 20, 29, 30, 45, 46, 47, 59]. These applications are advantageous since they have been shown to produce more biologically relevant results and take less computational time when compared to competing methods [16, 17, 46]. One of the advantages of utilizing non-procedural programming in biological applications is that biological knowledge can be naturally encoded into the program, instead of having to convert this information into procedural steps. Additionally, they restrict the search space of possible solutions and ensure only models that agree with constraint-encoded experimental data are retained [26]. It is expected that similar advantages will be achieved by applying non-procedural programming to predicting the 3D genomic organization. Here we present three mathematical modelling solutions to the 3D-GRP, collectively referred to as sonHi-C (pronounced “sonic”). Each mathematical model was implemented in a non-procedural language and tested with a normalized whole-genome contact map from fission yeast (Gene Expression Omnibus accession number: GSM1379427) [42]. Fission yeast was selected since it is a well-studied model organism with a relatively small but complex genome [58]. Additionally, it has many properties governing genomic organization that have been previously established using a variety of microscopy techniques.

## 3 Results & Discussion

This section presents the results for three mathematical modelling solutions to the 3D-GRP (Subsection 3.1), the non-procedural implementation for each mathematical model (Subsection 3.2) and a visualization of the predicted genome model (Subsection 3.3). Subsection 3.4 demonstrates the effect of varying the *m* parameter (described below) for the fission yeast dataset and Subsection 3.5 discusses how this research could be applied to organisms with higher ploidies (the number of chromosome copies) and/or larger genomes. All of the auxiliary files and programs used to generate these results are available at the project home page (https://github.com/kimmackay/SonHi-C).

### 3.1 Mathematical Modelling

Under normal cellular conditions, a given genomic region can be simultaneously involved in more than one interaction within the genome [18]. In contrast, a single genomic region within an individual cell is only able to participate in one Hi-C mediated interaction due to inherent restrictions within the biochemistry of the Hi-C experimental protocol [56]. In diploid organisms (organisms with two genomic copies) single cell Hi-C reactions are only able to detect two Hi-C mediated interactions per genomic region, one for each genomic copy [43]. An analogous restriction can be assumed in haploid organisms (organisms with only one genomic copy), where a single genomic region can only be actively detected in one Hi-C mediated interaction in a single cell. Using this restriction, a model of the 3D genome organization can be constructed from a whole-genome contact map by selecting a ploidy-dependent subset of the interactions for each genomic region that maximizes the sum of the corresponding interaction frequencies. The mathematical models and programs presented in this paper focus on modelling the 3D organization of haploid genomes but, as outlined in Subsection 3.5, they could be easily extended to organisms with higher ploidies.

Naively, a greedy heuristic could be employed to model the 3D fission yeast genome organization using the strategy described above. Briefly, the subset of interactions representing the solution set would be chosen by sorting and selecting the interactions with the largest corresponding frequency values. This process would then be repeated, rejecting any frequency that involves a region of the genome that has already been selected. This heuristic will fail to take into account the situation where lower frequencies, which were rejected by selecting a higher frequency interaction, actually result in a greater overall maximum value for the sum of all selected frequencies within the solution. An example of this can be seen in Figure 3 where panel A is a hypothetical whole-genome contact map and panels B and C represent two possible solution matrices with different overall frequency sums. Specifically, Figure 3B follows the greedy heuristic described above which results in a non-optimal solution where the selected frequencies sum to 1.3. Figure 3C shows the optimal solution where the selected frequencies sum to 1.4. This type of optimization problem has been shown to be well-suited for non-procedural approaches.

**Figure 3:**
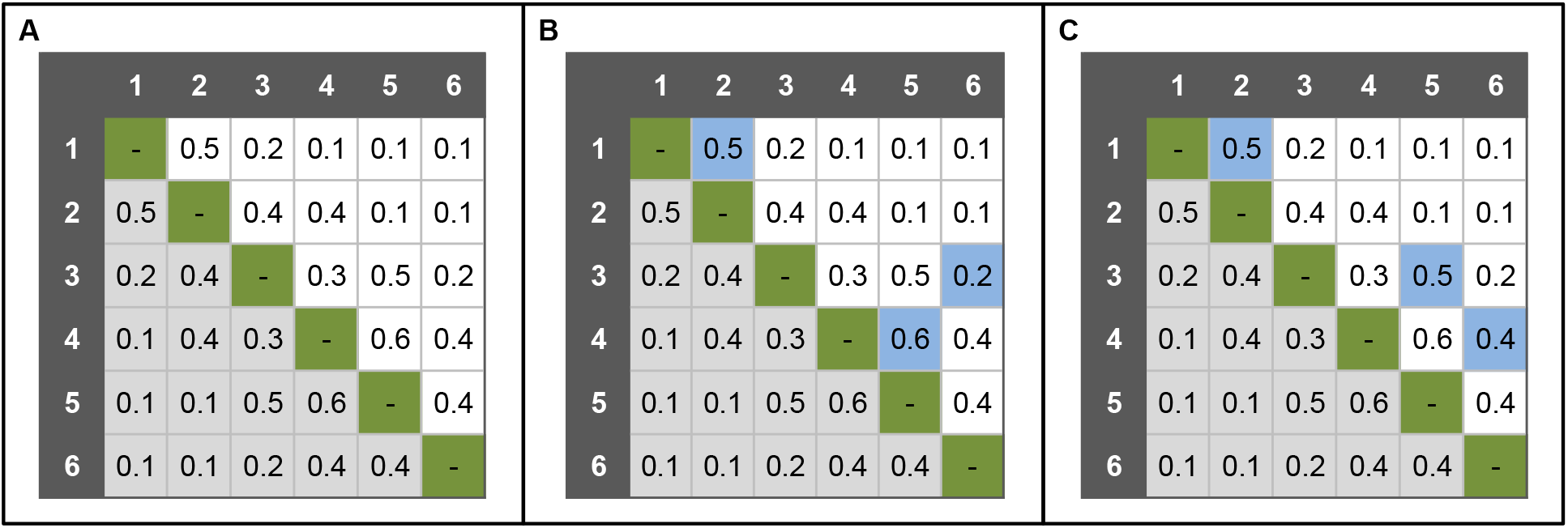
An example of two (of many) possible solutions to a 3D genome reconstruction problem. For all of the panels: the symmetric lower half of the contact map is indicated in light grey, the diagonal that represents “self-self” interactions is indicated in green and the genomic bin labels are represented in dark grey. For panels B and C: the blue boxes represent the subset of frequencies that could be selected as possible solutions (for *m* =1). Panel B is a representation of a valid, non-optimal solution from the greedy algorithm and panel C is a representation of the valid optimal solution for the contact map where the sum of the selected interaction frequencies are 1.3 and 1.4, respectively.

We developed three mathematical solutions to the 3D-GRP which describe the relationships present within the whole-genome contact map. As mentioned previously, a whole-genome contact map is a *N × N* matrix where the genome has been partitioned into *N* genomic bins. For a hypothetical whole-genome contact map (**A**), each cell (*A_i,j_*) records the normalized interaction frequency between genomic bins *i* and *j*. By construction, the contact map is symmetric (*A_i,j_* = *A_j,i_* for all *i, j*), and its main diagonal elements are all zero (*A_i,i_* = 0 for all *i*). The second parameter of our mathematical models is the maximal number of interactions that a given genomic bin can be involved in based on the source organism’s ploidy (denoted by the parameter *m*). For instance, *m* would be set to the following values based on the number of chromosome copies present: *m* =1 (haploid), *m* =2 (diploid; common in mammals), *m* = 4 (tetraploid; common in plants), and so on. Below are the specifics for three mathematical formulations of the 3D-GRP based on a given whole-genome contact map (**A**). Some variants require that the interaction frequencies be rounded and scaled to integer values.

#### 3.1.1 Constraint Programming (CP)

Our first model, **CP**, is encoded with Constraint Programming [49]. This model is valid for *m* = 1 only and requires integral *A_i,j_* values due to the implementation (described in Subsection 3.2.2). It is based on introducing variables *M_i_* where *M_i_* = *j* if genomic bin *i* interacts with genomic bin *j*, and *M_i_* = *i* otherwise. The goal of this model is to solve *M_i_* for all *i*. The model is given in Mathematical Model 1. Since this model encodes a combinatorial problem, its time complexity is exponential in the worst case.

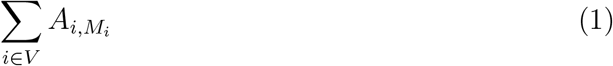

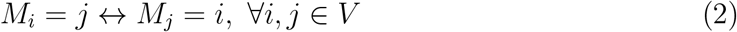

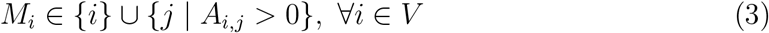

Mathematical Model 1: The **CP** model, valid for *m* =1 and integral interaction frequencies (*A_i,j_*) only. *V* is the set {1,…, *N*} representing the genomic bins.

#### 3.1.2 Graph Matching (GM)

Our second model, **GM**, is only valid for the *m* = 1 case. By representing the contact map as an undirected graph with *N* vertices (genomic bins) and 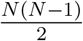 edges (interactions) the 3D-GRP can be regarded as the problem of computing a *maximum-weight matching* for the graph *G* = (*V, E*). A *matching* in a graph is a set of edges where no two edges share an endpoint. Each edge has an associated weight, and the weight of the matching is simply the sum of the weights of the edges in the matching. In the **GM** model, the vertices *V* are the set of genomic bins, the edges *E* are the set {(*i, j*) | *i* < *j* ^ *A_i,j_* > 0}, and the weights are given by *A*. An *O*(|*V*| · |*E*| log |*V*|) implementation of the weighted matching problem was reported by Mehlhorn and Schäfer [38], and is provided in the LEDA algorithm library ^1^. However, this formulation does not guarantee that each vertex in the original graph is represented in the matching. In terms of the 3D-GRP, this means that there is no guarantee each genomic bin from the contact map would be represented in the solution. In order to overcome this, the 3D-GRP can be represented as a maximum-weight *perfect* matching problem to ensure *all* vertices in the graph are matched. Edmonds [15] invented the first polynomial maximum-weight perfect matching algorithm, which runs in *O*(|*V*|^2^|*E*|) time. If the graph was completely connected (i.e. 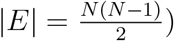, this would intuitively suggest a time complexity of *O*(*N*^4^), but in reality whole-genome contact maps are characteristically sparse resulting in |*E*| ≪ *N*^2^ since zero-weight edges are not represented in the graph. As such, the mean computational complexity will depend on the experimental resolution and resultant sparsity of a given whole-genome contact map. Kolmogorov’s Blossom V algorithm [28] is considered the most efficient implementation of Edmonds algorithm. We use this reduction in our model, given in Mathematical Model 2.

Solve the maximum-weight perfect matching problem for the graph *G′* = (*V′, E′*) and weight function 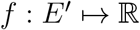, i.e.:

maximize

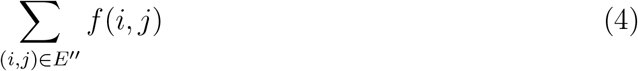

subject to:

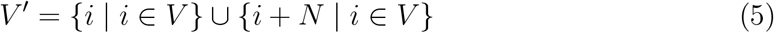

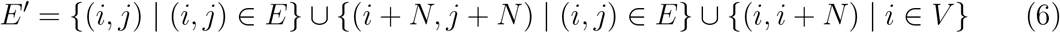

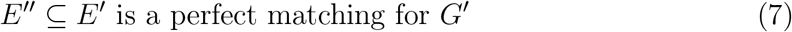

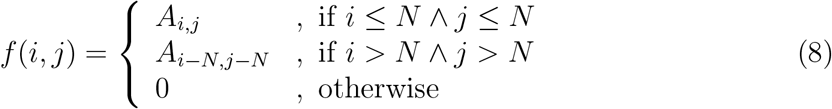

Mathematical Model 2: The **GM** model, for *m* = 1 only. *V* is the set {1, …, *N*} representing the genomic bins. *E* is the set {(*i, j*) | *i* < *j* ^ *A_i,j_* > 0} representing the interactions and the weights are given by *A*. *f*(*i,j*) is the function used to calculate edge weight. *G* = (*V′*, *E′*) is an extended graph used to map *G* = (*V, E*) to a maximum-weight perfect matching problem. This mapping to maximum-weight perfect matching was given by Mehlhorn [38, footnote 1].

#### 3.1.3 Integer Programming (IP)

Our third model, **IP**, uses Integer Programming [57] and is valid for any value of *m*. It is based on introducing variables *x_i,j_* that assume a value of 1 if genomic bin *i* interacts with genomic bin _j_, and 0 otherwise. The goal of this model is to solve *x_i,j_* for all *i, j*. The model is given in Mathematical Model 3. Similarly to the **CP** model, this model encodes a combinatorial problem resulting in an exponential worst-case time complexity.

maximize

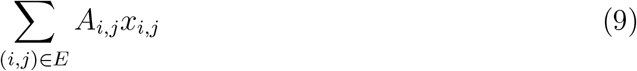

subject to:

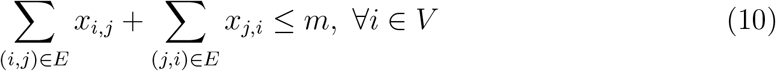

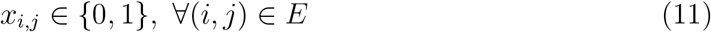

Mathematical Model 3: The **IP** model, for any *m*. *V* is the set {1,…, *N*} representing the genomic bins. *E* is the set {(*i, j*) | *i* < *j* ∧ *A_ij_* > 0} representing the interactions and the weights are given by *A*.

### 3.2 Implementations

Each mathematical model described above was implemented in a non-procedural language (specifics are provided below) and tested with an existing fission yeast Hi-C dataset (GEO accession number: GSM1379427 [42]). These implementations were run on a server-grade computer with sufficient main memory to represent the entire problem. When the implementations were run on the complete fission yeast whole-genome contact map (results presented in Subsections 3.2.3 and 3.2.4), there were only a few *trans*-chromosomal interactions within the solution sets making it difficult to infer the organization of the chromosomes in relation to each other. The low number of *trans*-chromosomal interactions is likely due to the fact that *cis*-chromosomal interactions are known to have higher interaction frequencies than *trans*-chromosomal interactions within the genome [12, 33]. This makes it more likely for *cis*-chromosomal interactions to be included in the solution set since the goal of the mathematical models described above is to select a maximal subset of interaction frequencies. To overcome this, a divide and conquer approach was used where each *cis*-chromosomal and pairwise *trans*-chromosomal subproblem was locally solved. These solutions were then merged to retain the selected *cis*- and *trans*-chromosomal interactions from each subproblem (described in Subsection 3.2.1). Overall, this divide and conquer approach resulted in a larger number of trans-chromosomal interactions being included in the final solution set.

#### 3.2.1 Divide & Conquer

As mentioned above, the *cis*-chromosomal subproblems better represent the individual chromosome structure while the *trans*-chromosomal subproblems represent how the chromosomes are organized in relation to each other within the nucleus. In the case of the fission yeast dataset six separate subproblems were generated: one for each chromosome’s *cis*-interactions and one for each set of pairwise *trans*-chromosomal interactions (this division heuristic is described below). Each subproblem was independently run and the results were merged to generate the visualizations presented in Subsection 3.3. This is one of the first times a divide-and-conquer approach has been applied to 3D genome prediction from Hi-C data (cf. [48]).

##### Divide

A single whole-genome contact map can be naturally divided into a finite, organism specific number of subproblems representing its constituent *cis*-interactions and pairwise *trans*-interactions. Each subproblem can be defined within the whole-genome contact map by specifying the range of genomic bins that correspond to the *cis-* or *trans*-interactions for each chromosome. In general, the number of subproblems for a whole-genome contact map with *k* chromosomes is equal to 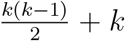 where 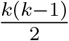 represents the number of pairwise *trans*-interaction subproblems and *k* represents the number of *cis*-interaction subproblems. For example, because fission yeast has three chromosomes, its whole-genome contact map can be naturally partitioned into six subproblems (three *cis*- and three *trans*-interaction subproblems) to be solved in parallel. The location of these subproblems within a fission-yeast whole-genome contact map are depicted in Figure 4.

**Figure 4:**
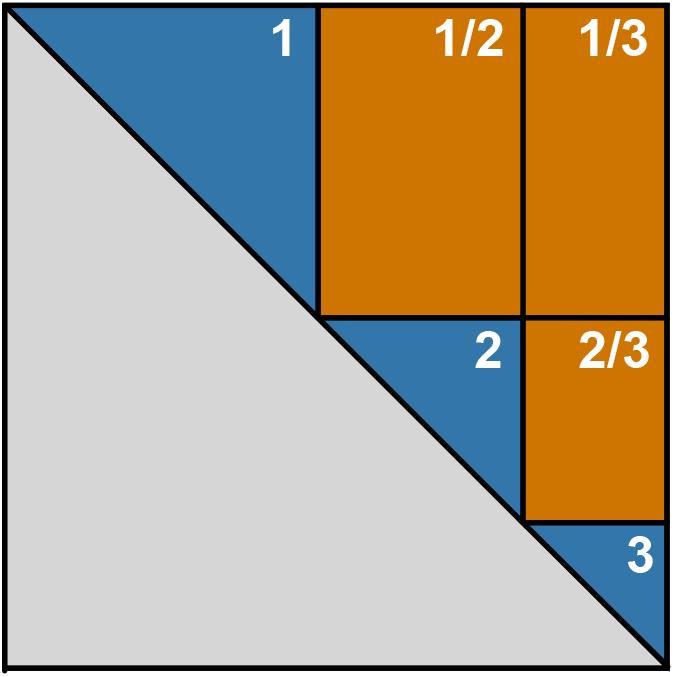
Identification of subproblems within the fission yeast contact map. The large grey triangle represents the portion of the contact map that does not need to be processed since all contact maps are mirrored along the diagonal. The blue triangles represent the subsections of the contact map that correspond to intra-chromosomal interactions, while the orange squares represent the subsections of the contact map that correspond to the inter-chromosomal interactions. The labels on the blue and orange areas represent the chromosome(s) involved in the interactions within that subsection of the contact map. In terms of the intra-chromosomal interactions, chromosome 1 contains the largest number of genomic bins while chromosomes 2 and 3 account for 34 and 80 percent fewer bins, respectively.

##### Local Conquer

In order to solve the entire 3D-GRP, programs corresponding to the *cis*-interaction subproblems and pairwise *trans*-interaction subproblems can be generated and run independently. The results can then be combined using the merge step described below.

##### Merge

The solutions from each subproblem are combined to reconstruct the entire 3D genomic model. This step is a heuristic which utilizes a novel coefficient (called the “dynamics coefficient”) to account for the instances when a single genomic region participates in more than *m* subproblem solutions; i.e. more than *m* interactions. Instead of eliminating interactions from subproblem solutions involving the same genomic region (when this region has already been selected in *m* interactions), each identified interaction is maintained and associated with a region-specific dynamics coefficient to encode the mobility (or lack of mobility) of that genomic region. Briefly, the dynamics coefficient for each genomic region is calculated by scanning all of the resultant files for each subproblem and counting how many times a specific genomic bin is found across the subproblem solution sets. The more interactions a genomic region is involved in, the higher its corresponding dynamics coefficient, and *vice versa*, In general, the dynamics coefficient is an integer value in the range of 0 to *k* where *k* is the number of chromosomes present in the genome. For example, in fission yeast (*k* = 3) if genomic bin 1 was involved in an interaction in the solution sets of the chromosome 1 *cis*-interaction subproblem and the chromosome 1/2 *trans*-interaction subproblem it would have a dynamics coefficient of 2, whereas if it was involved in an interaction in each of the relevant *trans*-interaction subproblems and the *cis*-interaction subproblem it would have an associated dynamics coefficient of 3. A higher dynamics coefficient suggests that the corresponding genomic region was more mobile within the genome and that there was less certainty about its fixed position within the model. This is similar to the B factor (also known as the temperature factor or the Debye-Waller factor) generated with protein X-Ray Crystallography experiments [32]. The B factor encodes the degree of uncertainty associated with computed atomic positions in 3D space.

Utilizing the dynamics coefficient allows for the overall solution to the 3D-GRP to retain the information associated with each subproblem’s optimal solution instead of having to exclude interactions that involve genomic regions already selected *m* times as part of the solution set. Although this violates the initial ploidy restriction used to constrain the mathematical models, it is still biologically valid. As mentioned previously, it is possible for a given genomic bin to be involved in more than one interaction in 3D space [8, 18], even though Hi-C is only able to detect one pairwise interaction per restriction site within a single haploid cell. Additionally, the dynamics coefficient allows the program to encode some of the mobility of genome organization into the predicted model by representing the certainty of whether an interaction is fixed within the population of cells. Finally, the dynamics coefficient is also used to calculate relative distances between genomic bins which are used to visualize the predicted model.

#### 3.2.2 CP model

The **CP** mathematical model (depicted in Mathematical Model 1) was implemented in MiniZinc [44] with the OR-Tools constraint solver from Google^2^. An example MiniZinc program (Program 1) and a corresponding example data file (Example Data File 4) with the integral interaction frequencies from the hypothetical whole-genome contact map depicted in Figure 3A are given in the Appendix. This model leverages the fact that the solution will never contain more than *m* × *N* interactions making it scalable to larger genomes in terms of space complexity. It is worth noting that Equation (2) can be encoded by the inverse global constraint^3^, whereas Equation (1) is encoded with one element constraint per row of A plus one sum constraint. These constraints are propagated by efficient algorithms in many constraint programming solvers.

The MiniZinc program corresponding to the complete fission yeast genome could not be solved to optimality after several days of run time on a server-grade computer. In an attempt to overcome this, the divide-and-conquer approach described above was applied. A MiniZinc program for each *cis*- or *trans*- subproblem was generated and run independently. Similarly to the complete whole-genome contact map, not a single *cis*- or *trans*- problem could be solved to optimality in several days.

#### 3.2.3 GM Model

The **GM** mathematical model (depicted in Mathematical Model 2) was implemented in SICStus Prolog^4^ [6] using Kolmogorov’s Blossom V algorithm [28]. The implemented program using this representation is presented in Program 2. An example associated data file for this program is given in Example Data File 5 and is based on the interaction frequency values from the hypothetical whole-genome contact map depicted in Figure 3A. The program is run by: (1) invoking the `compile_adjacency` predicate with a data file similar to that given in Example Data File 5 and (2) invoking the `match_blossom5` predicate. In this example, this would be done by invoking: `compile_adjacency (‘testMap.csv’, testMap)`, followed by `match_blossom5(testMap,[1],[1])`. For the fission yeast results, all of this has been automated in a “makefile” that is available on the project homepage (https://github.com/kimmackay/SonHi-C).

The SICStus Prolog implementation of the **GM** mathematical model was able to predict a fission yeast genomic organization in 1.088 seconds (*m* = 1; for the complete whole-genome contact map where |*V*| = 1258, |*E*| = 745595). In this matching, only one edge representing a *trans*-chromosomal interaction was included while the rest of the edges depicted *cis*-chromosomal interactions. This made it difficult to infer the organization of the chromosomes in relation to each other. In order to overcome this the divide-and-conquer approach described above was applied. Specifically, six separate matchings were identified: one for each chromosome’s *cis*-interactions and one for each set of pairwise *trans*-chromosomal interactions. A SICStus Prolog program for each *cis*- or *trans*-subproblem was run independently. For each subproblem, the time it took to identify the optimal solution is presented in Table 1. These results were merged using the `generate_gephi_input_subproblems.pl` script available at the project homepage. The merge step took less than 1 second of execution time.

**Table 1.** Subproblem sizes and corresponding run times for the **GM** mathematical model applied to the fission yeast whole-genome contact map.

| Subproblem | Number of Vertices ($\vert V\vert $) | Number of Edges ($\vert E\vert $) | Local Conquer Runtime (seconds) |
| --- | --- | --- | --- |
| chromosome 1<br>*cis*-interaction | 558 | 148734 | 0.164 |
| chromosome 2<br>*cis*-interaction | 454 | 96562 | 0.055 |
| chromosome 3<br>*cis*-interaction | 246 | 27255 | 0.008 |
| chromosome 1/2<br>*trans*-interaction | 454 | 241022 | 9.645 |
| chromosome 1/3<br>*trans*-interaction | 246 | 128472 | 7.203 |
| chromosome 2/3<br>*trans*-interaction | 246 | 103550 | 1.552 |

#### 3.2.4 IP Model

The **IP** mathematical model (depicted in Mathematical Model 3) was implemented in Prolog and solved using the mixed integer programming based Gurobi Optimizer ^5^ [25]. The implemented program using this representation with the hypothetical whole-genome contact map depicted in Figure 3A is shown in Program 3. This implementation uses the same data file as the **GM** model (Example Data File 5). The program is run by: (1) invoking the `compile_adjacency` predicate with a data file similar to that given in Example Data File 5 and (2) invoking the `solve_ip` predicate. For this example, this would be done by invoking: `compile_adjacency(‘testMap.csv’, testMap)`, followed by `solve_ip(testMap,[1],[1])`. For the fission yeast results, all of this has been automated in a makefile that is available on the project homepage (https://github.com/kimmackay/SonHi-C).

The Prolog program for the complete whole-genome fission yeast contact map was able to predict a genomic organization in 294.44 seconds (*m* = 1; |*V*| = 1258, |*E*| = 745595). Similarly to the **GM** models’ solution, only one *trans*-chromosomal interaction was represented in the solution set. The same trans-chromosomal interaction was present in the **GM** and **IP** solutions. The presence of only one *trans*-chromosomal interaction made it difficult to infer the organization of the chromosomes in relation to each other. In order to overcome this, the divide-and-conquer approach described above was used. Six separate subprograms were generated and run independently (one for each chromosome’s *cis*-interactions and one for each set of pairwise *trans*-chromosomal interactions). The size of each problem in terms of *V* and *E* as well as the time it took to identify the optimal solution is presented in Table 2. In each case, the optimal solution identified was identical to the matching reported by the **GM** model. These results were merged using the `generate_gephi_input_subproblems.pl` script available at the project homepage. The merge step took less than 1 second of execution time.

**Table 2.** Subproblem sizes and corresponding run times for the **IP** mathematical model applied to the fission yeast whole-genome contact map.

| Subproblem | Number of Vertices ($\vert V\vert $) | Number of Edges ($\vert E\vert $) | Local Conquer Runtime (seconds) |
| --- | --- | --- | --- |
| chromosome 1<br>*cis*-interaction | 558 | 148734 | 15.75 |
| chromosome 2<br>*cis*-interaction | 454 | 96562 | 9.26 |
| chromosome 3<br>*cis*-interaction | 246 | 27255 | 2.06 |
| chromosome 1/2<br>*trans*-interaction | 454 | 241022 | 4.90 |
| chromosome 1/3<br>*trans*-interaction | 246 | 128472 | 4.70 |
| chromosome 2/3<br>*trans*-interaction | 246 | 103550 | 3.40 |

### 3.3 Visualization

Since the results for the **GM** and **IP** models were identical, only one visualization is shown here. The results were converted into an undirected graph and visualized using Gephi (Figure 5A) [53, 54]. These images are graph-based visualizations of the predicted model based on the graphical representation of Hi-C data described in GrapHi-C [37]. Briefly, the nodes in the network represent the individual genomic bins of the whole-genome contact map and the edges represent either selected interactions between bins or known linear interactions between adjacent bins. Linear interactions add additional biological constraints by representing the *bonafide in vivo* linear connections between bins (i.e. the linear extent of the chromosome). Each edge was weighted using either: the interaction frequency divided by the dynamics coefficient (for *cis*- and *trans*- interactions) or the experimental resolution (for linear interactions). The Force Atlas 2 layout was then applied to the network and the nodes were coloured according to their chromosome number or genomic feature. We would like to stress that this graph-based visualization is not a polymer model of the DNA chain that is often seen in other 3D genome prediction tools. Therefore, the smoothness of the edges is not a result of any bending rigidity constraints. Instead, it is a result of the visualization tool (Gephi) and the network layout applied.

**Figure 5:**
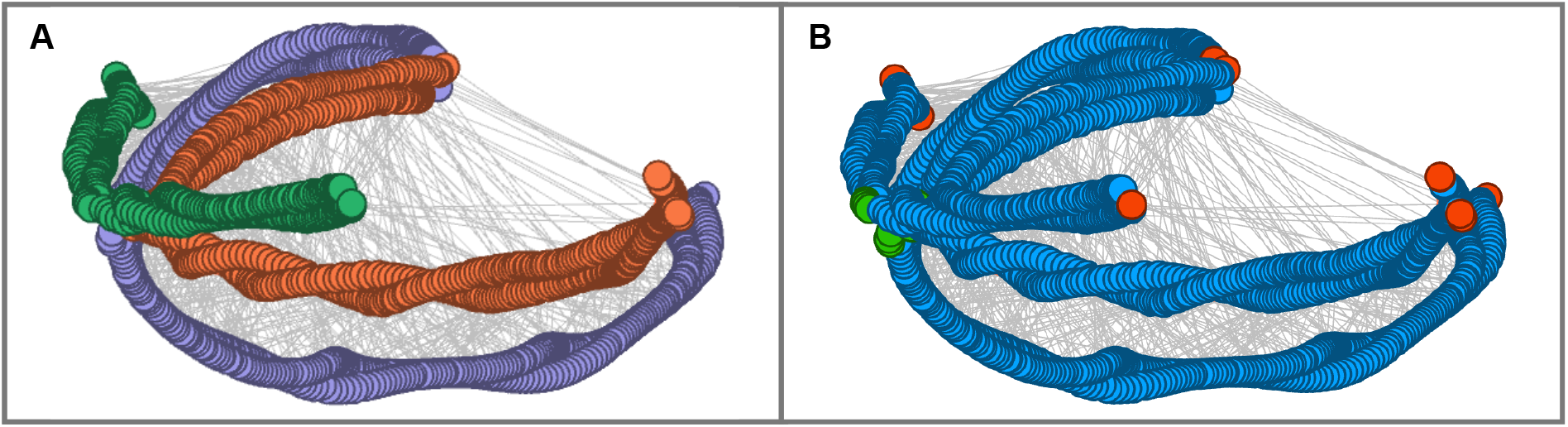
Visualization of the Predicted Genome Model Using the **GM**/**IP** Models and Identification of Genomic Features Indicative of Fission Yeast Mitotic Chromosomal Organization. Circles depict the genomic bins, grey lines represent *cis*- and *trans*-interaction edges selected by the **GM**/**IP** models, and line lengths are proportional to the associated dynamics coefficient and the inverse of the interaction frequency (*A_i,j_*). In Panel A, circles are coloured according to their corresponding chromosome (CHR1: purple, CHR2: orange, CHR3: green). In Panel B, the following genomic features are highlighted: telomeres (red), centromeres (green) and nuclear DNA (blue).

One of the most well–documented features of fission yeast genomic organization is the 3D clustering of centromeres and telomeres within the nucleus [9, 19]. In order to determine whether the predicted yeast model was able to recapitulate these features, the genomic bins corresponding to centromeres and telomeres were coloured in the Gephi visualization. Figure 5B provides a visual depiction of the location of the centromeres and telomeres in the predicted genomic model. This figure provides further evidence that the predicted genome model is consistent with established principles of mitotic fission yeast chromosomal organization including: (1) chromosomal organization into a hemispherical region, (2) a single centromere cluster and (3) the presence of two telomere clusters (chromosome 1/2) located near the nuclear periphery, opposite the centromere cluster [55]. The clustering of the regions appears to be conserved in the predicted model providing confidence in the biological accuracy that was achieved using the **GM** and **IP** mathematical models and the corresponding non-procedural implementations.

### 3.4 Effect of *m* on Genome Organization in Fission Yeast

As mentioned previously, it is possible that each genomic region could be involved with more than one interaction within the genome but is restricted to *m* Hi-C interactions (where the value of *m* is based on organism ploidy). To determine whether or not relaxing this ploidy restriction would result in a more comprehensive genomic model, the implemented program for the **IP** model was tested with values of *m* from 1 to 6 for the same fission yeast Hi-C dataset used above (GSM1379427 [42]). As mentioned previously, this mathematical model allows for a single genomic bin to be involved in more than one Hi-C mediated interaction in the predicted genome organization. For each value of *m*, the program was able to find an optimal solution in 294.44 seconds, 13.20 seconds, 104.46 seconds, 15.31 seconds, 38.79 seconds, and 16.94 seconds for *m* = 1‥6, respectively. Similarly to what was described above, the results for each value of *m* were converted into a graph and visualized using Gephi (Figure 6). Each edge was weighted using either: the interaction frequency divided by the dynamics coefficient (for *cis*- and *trans*- interactions) or the experimental resolution (for linear interactions). The Force Atlas 2 layout was then applied to the network and the nodes were coloured according to their chromosome number. The nodes in the graph represent the individual genomic bins of the whole-genome contact map and the edges represent either selected interactions between bins or known linear interactions between adjacent bins.

**Figure 6:**
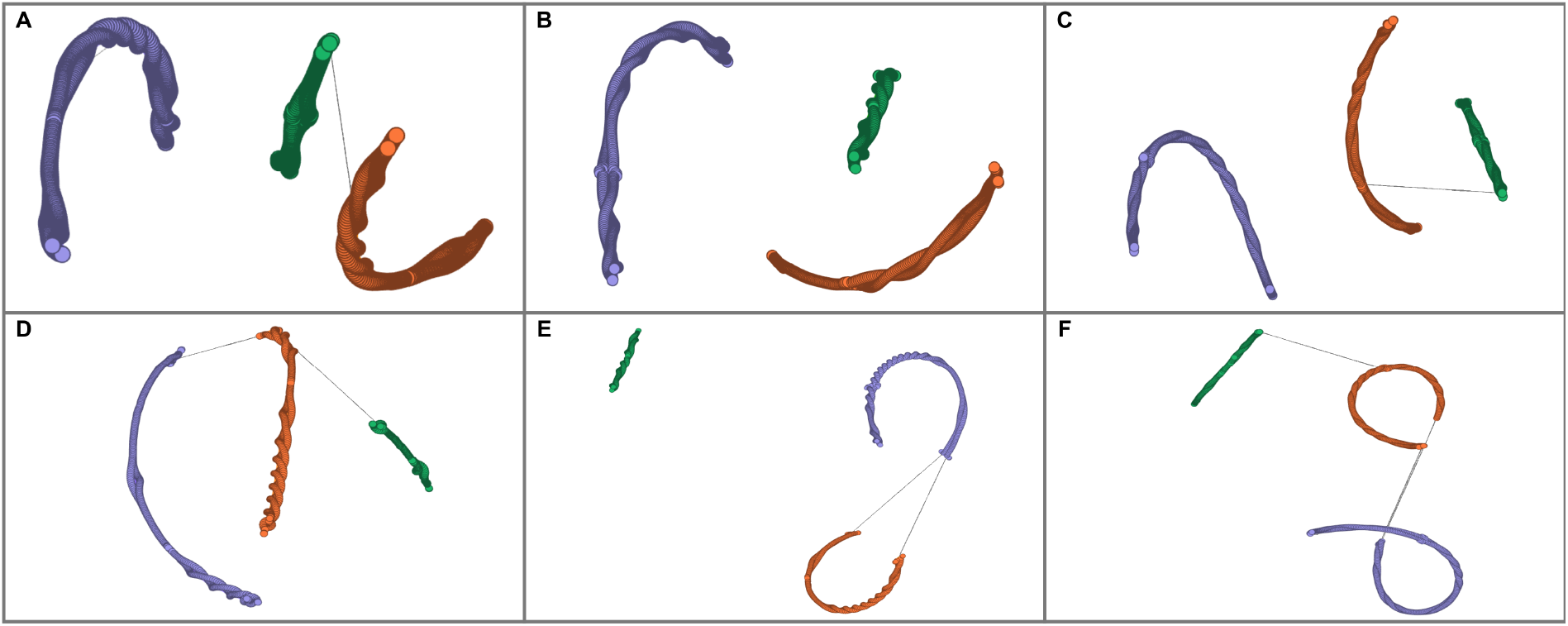
Visualization of the Predicted Genome Model Using the **IP** Model with Various *m* Values. Circles depict the genomic bins, grey lines represent *trans*-interaction edges selected by the **IP** model, and line lengths are proportional to the associated interaction frequency (*A_i,j_*). Circles are coloured according to their corresponding chromosome (CHR1: purple, CHR2: orange, CHR3: green). The results for each *m* values are presented in the following panels: A (*m* = 1), B (*m* = 2), C (*m* = 3), D (*m* = 4), E (*m* = 5), F (*m* = 6).

The results presented in Figure 6 indicate that relaxing the ploidy restriction (by increasing the value of *m*) does not result in a more comprehensive genomic model. Similarly to the **GM** and **IP** whole-genome predictions, minimal *trans*-chromosomal interactions were selected by the model regardless of what the parameter *m* was set to. Specifically the following number of *trans*-chromosomal interactions were observed in each solution set: 1 (*m* = 1), 0 (*m* = 2), 1 (*m* = 3), 2 (*m* = 4), 2 (*m* = 5), 3 (*m* = 6). It is clear that regardless of the number of interactions allowed per genomic bin, separating the 3D-GRP into *cis*- and *trans*subproblems is a more viable strategy for predicting genome organization when using the **IP** mathematical model. This is likely due to the fact that *cis*-chromosomal interactions occur more frequently than *trans*-chromosomal interactions within the genome (resulting in higher interaction frequency values) [12].

### 3.5 Application to Organisms with Higher Ploidies and/or Larger Genomes

The **IP** mathematical model described above could be easily applied to organisms with higher ploidies. As mentioned previously, the *m* parameter defines the number of Hi-C mediated interactions in which each genomic bin can actively be participating within the solution set. Therefore, this model could be easily applied to organisms with higher ploidies by specifying the value of the *m* parameter. For instance, *m* could be modified in the following ways according to the number of chromosome copies present: *m* = 2 (diploid; common in mammals), *m* = 4 (tetraploid; common in plants), and so on. One issue that would need to be addressed in organisms with higher ploidies is phasing the interactions to each chromosome copy. This could potentially be solved using existing phasing tools [10] and additional biological data [4, 51].

Utilizing the divide-and-conquer approach described in Section 3.2 allows one to take advantage of coarse-grained parallelism ensuring the mathematical models are scaleable to organisms with larger genomes (for *m* ≥ 1). This type of parallelism is easy to obtain on many types of computational infrastructure. As an example, it could be easily applied to a whole-genome contact map from *Homo sapiens*. This contact map would result in the generation of 276 subproblems (given *k* = 23 and the number of subproblems 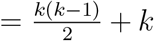). It would not be hard to find 276 cores in the current environment of computer grids and multi-core machines to run the problems representing the subproblems in parallel. Since each subproblem can be run in parallel and the merge step has a linear time complexity (in terms of the number of genomic bins), the associated average runtime of the computation is expected to be polynomial in the number of genomic bins. In general the complexity will depend on the chromosome size and the associated experimental resolution).

## 4 Future Work

Future work will focus on the validation, modification and extension of the 3D-GRP solutions presented in this manuscript. Specifically, an extensive biological validation of the predicted genome models will be performed to better characterize the biological accuracy of the developed mathematical models. The formulation for the **CP** mathematical model will be modified to adjust the proportion of *cis*- and *trans*-chromosomal interactions in the solution set. Additionally, different types of data transformations will be investigated to better accommodate the proportion of *cis*- and *trans*-chromosomal interactions in the whole-genome contact map. The **GM** model will be parameterized so that it can be used with different values of *m*. The **IP** mathematical model will be utilized as a computational framework which will be extended and further developed to incorporate a variety of additional genomic datasets and information types into the prediction process. The non-procedural representation utilized in this computational framework will allow additional datasets to be naturally incorporated into the prediction of the 3D genome. For example, each genomic bin could have an associated list of variables representing the genes found within that bin and their corresponding gene expression values. Constraints could then be applied to favour interactions between regions with similar expression profiles. The **IP** mathematical model will also be utilized as a starting point for predicting the 3D genomic structure of organisms with higher ploidies by applying the modifications suggested in the subsection above. Finally, the **IP** model could be adapted and used for detecting topologically associated domains [13, 22] within a whole-genome contact map.

## 5 Conclusion

This is the first time a non-procedural programming approach has been used to model the 3D genome organization from Hi-C data. Specifically, we developed three novel mathematical models for predicting the 3D genome from Hi-C data. Each mathematical model was implemented in a non-procedural language and tested with an existing Hi-C dataset from fission yeast. The **CP** model was not able to solve the 3D-GRP to optimality after several days of execution. The **GM** model was able to optimally solve the 3D-GRP for fission yeast in 1.088 seconds. The **IP** model was able to optimally solve the 3D-GRP in fission yeast for *m* =1 in 294.44 seconds resulting in the same solution as the **GM** model. In this solution, only one *trans*-chromosomal interaction was selected making it difficult to infer the organization of the chromosomes in relation to each other. A divide-and-conquer approach was used to overcome this where six separate subproblems (one for each set of *cis*-chromosomal interactions and one for each set of pairwise *trans*-chromosomal interactions) were independently solved, combined and visualized. The *cis*-chromosomal matchings better represent individual chromosome structure while the *trans*-chromosomal matchings represent how the chromosomes are organized in relation to each other within the nucleus. In the case of the **GM** and **IP** models, the predicted genome organizations represent the interactions of a population-averaged consensus structure where the most likely interactions are present (this is determined by maximizing the sum of the interaction frequencies of the selected interactions). Each predicted genome organization was then biologically validated through literature search which verified that the prediction recapitulated key documented features of the yeast genome. The divide-and-conquer solution strategy lends itself to additional speed improvements due to the potential for running the defined subproblems in parallel. Furthermore, a novel coefficient was defined (the dynamics coefficient) which allowed a level of positional uncertainty to be encoded into the predicted genomic organization. Overall, the mathematical models and programs developed here demonstrate the power of non-procedural applications for modelling the 3D genome and are a step towards a better understanding of the relationship between genomic structure and function.

## Declarations

### Ethics approval and consent to participate

Not applicable

### Consent for publication

Not applicable.

### Availability of data and material

The datasets supporting the conclusions of this article are available in the Gene Expression Omnibus database(accession number: GSM1379427; https://www.ncbi.nlm.nih.gov/geo/query/acc.cgi?acc=GSM1379427).

### Software information

Project Name: SonHi-C (pronounced “sonic”)

Project home page: https://github.com/kimmackay/SonHi-C

License: This work is licensed under the Creative Commons Attribution-NonCommercial-ShareAlike 3.0 Unported License. To view a copy of this license, visit http://creativecommons.org/licenses/by-nc-sa/3.0/ or send a letter to Creative Commons, PO Box 1866, Mountain View, CA 94042, USA.

## Author′s contributions

KM, MC, AK developed the **CP** and **GM** mathematical models. MC developed the **IP** mathematical model. MC implemented and tested the constraint models. KM visualized the results and verified the biological accuracy. KM wrote the manuscript. MC, AK edited the manuscript.

## Acknowledgements

We could like to thank Dr. Christopher Eskiw and Conor Lazarou for their input and advice.

## Authors’ information

The e-mail address for each author is as follows: Kimberly MacKay; Mats Carlsson; Anthony Kusalik.

## A Implemented Programs

Program 1: The MiniZinc implementation of the **CP** mathematical model.

~~~
1 %% Load the relevant libraries
2 include “globals.mzn”;
3
3 %% Variable Declarations
4 int: N;
6
5 %% first chromosome of interest
6 set of 1‥N: Chr1 ;
9
7 %% second chromosome of interest
8 set of 1‥ N: Chr2 ;
12
9 %% the given frequency map, assumed symmetric, main diagonal = 0
10 array [1‥N,1‥ N] of int: map;
15
11 %% main decision variables
12 array [1‥ N] of var 1‥ N: match ;
18
13 %% objective per row
14 array [1‥N] of var int: rowobj ;
21
15 %% total objective, N.B. counting each binding twice
16 var int : dobj ;
24
17 %% mask out all entries not connecting Chr1 and Chr2
18 int: masked_map(1‥N: i, 1‥N: j) =
19   if i in Chr1 /\ j in Chr2 then
20      map [i,j]
21   else if i in Chr2 /\ j in Chr1 then
22      map [i,j]
23   else 0 endif endif;
32
24 %% assertion: frequency map is symmetric
25 constraint
26   forall(i in 1‥N, j in 1‥N where i<j)
27     (assert(masked_map(i,j) = masked_map(j,i), “Asymmetry!”));
37
28 %% constrain the objective, one slice per row
29 constraint
30   forall(i in 1‥ N)
31     (rowobj [i] = [masked_map(i,j) | j in 1‥N][match[i]]);
42
32 %% constrain the total objective
33 constraint
34   dobj = sum(i in 1‥N)(rowobj [i]);
46
35 %% domination: prevent zero - frequency edges
36 constraint
37   forall (i in 1‥ N)
38     (match[i] in {i} union {j |j in 1‥N where masked_map(i,j)>    });
51
39 %% domination: prevent obviously suboptimal solutions
40 constraint
41   forall(i in 1‥N, j in 1‥N where
42     masked_map(i,j)>0)(rowobj [i] + rowobj[j] >= 1);
56
43 %% essential matching constraint
44 constraint
45   inverse(match, match) :: domain;
60
46 %% Solve
47 solve :: int_search(rowobj, max_regret, indomain_max, complete)
48   maximize (dobj);
64
49 %% output the results
50 output
51  [”edge (\(i),\(match[i])). % benefit = \(rowobj [i])\n” |
52   i in 1‥N where fix (match [i])>i] ++
53  [“ objective(\(dobj div 2)).\n”] ++
70  [] ;
~~~

Program 2: The implemented program using the **GM** mathematical model.

~~~
1 %% Note: this program assumes a local version of BlossumV
2 %% exists on the computer. The majority of this program formats the
3 %% input file in order to pass the data to BlossumV and parses
4 %% the output generated by that solver. The program makes use of the
5 %% temporary files /tmp/all.in and /tmp/all.out.
6 %% The call to BlossumV is highlighted in red.
7
7 :- use_module(library(lists)).
8 :- use_module(library(csv)).
9 :- use_module(library(system3)).
11
10 chromosome (testMap, 1, 1, 6).
13
11 genome_size (testMap, 6).
15
12 scale_factor(testMap, 1.0E1).
17
13 compile_adjacency(Path, Species) :-
14   retractall (adjacency(_,_,_,_,_)),
15   see(Path),
16   read_record (_),
17   repeat,
18     read_record (Record),
19     (Record = end_of_file -> true
20 ; parse_record(Record, B1, B2, F),
21   bin2chr(B1, Species, Chr1),
22   bin2chr(B2, Species, Chr2),
23   assertz(adjacency(B1, B2, F, Chr1, Chr2)),
24   fail
25 ), !,
26  seen,
27  save_predicates([adjacency/5], Species).
33
28 match_blossom5(Species, Set1, Set2) :-
29   load_files(Species),
30   retractall(edge(_,_,_)),
31   (adjacency(B1, B2, F, Chr1, Chr2),
32     B1 < B2,
33     (member(Chr1, Set1), member(Chr2, Set2) -> true
34     ; member(Chr1, Set2), member(Chr2, Set1) -> true
35     ),
36     assertz (edge (B1, B2, F)),
37     fail
38 ;   true
39  ),
40  findall(edge(I,J,W), edge(I,J,W), Edges),
41  length (Edges, E),
42  genome_size(Species, N),
43  scale_factor(Species, Scale),
44  tell(‘/tmp/all.in ‘ ),
45  NN is 2*N,
46  EEN is 2*E+N,
47  print_tab ([NN, EEN]),
48  (foreach (edge (I1,J1,W1), Edges ),
49    param(Scale)
50  do I11 is I1-1,
51     J11 is J1-1,
52     W11 is integer (-Scale*W1),
53     print_tab([I11,J11,W11])
54 ),
55 (foreach(edge(I2,J2,W2),Edges),
56     param (N, Scale )
57 do  I21 is I2+N-1,
58     J21 is J2+N-1,
59     W21 is integer(-Scale*W2),
60     print_tab ([I21, J21,W21] )
61 ),
62 (for (K,1,N),
63     param(N)
64 do  K1 is K-1,
65     K2 is K+N-1,
66     print_tab ([K1,K2,0] )
67 ),
68 told,
69 system(′blossom5 -e /tmp/all.in -w /tmp/all.out ‘),
70  see(‘/tmp/all.out ‘ ),
71  retractall(solution(_,_,_)),
72  read_line (_),
73  repeat,
74    read_line(Codes),
75    (Codes = end_of_file -> true
76     ; append(Icodes, [0 ‘ |Jcodes], Codes),
77     number_codes(I0, Icodes),
78     number_codes(J0, Jcodes),
79     I is I0 + 1,
80     J is J0 + 1,
81     edge (I, J, W),
82     assertz(solution(I,J,W)),
83     fail
84  ), !,
85  seen,
86  findall(W, solution(_,_,W), Ws),
87  length(Ws, Size),
88  sumlist(Ws, Weight),
89  format(‘ % Blossom V computed a matching of size ~d and weight ~w\n’,
90      [Size, Weight ]),
91  findall(0, (solution(I,J,W), print_tab([I,J,W])), _).
98
92 print_tab (L) :-
93   (foreach (X,L) do write(X), write(‘ ‘ )),
94    nl .
102
95 parse_record (Record, B1, B2, F) :-
96   Record = [integer(B1,_), integer(B2,_), float (F,_)].
105
97 bin2chr(Bin, Species, Chr) :-
98   chromosome (Species, Chr, A, B),
99   Bin >= A,
100  Bin =< B, !.
~~~

Program 3: The implemented program using the **IP** mathematical model.

~~~
1 %% Note: this program assumes a local version of the Gurobi solver
2 %% exists on the computer. The majority of this program formats the
3 %% input file in order to pass the data to Gurobi and parses
4 %% the output generated by that solver. The program makes use of the
5 %% temporary files /tmp/all.lp and /tmp/all.sol.
6 %% The call to Gurobi is highlighted in red.
7
7 :- use_module(library(lists)).
8 :- use_module(library(csv)).
9 :- use_module(library(system3)).
11
10 chromosome (testMap, 1, 1, 6).
13
11 solve_**IP** (Species, Set1, Set2, M) :-
12 load_files (Species ),
13 retractall (edge (_,_,_)),
14 (adjacency(B1, B2, F, Chr1, Chr2),
15   B1 < B2,
16 (member(Chr1, Set1), member(Chr2, Set2) -> true
17 ; member(Chr1, Set2), member(Chr2, Set1) -> true
18 ),
19 assertz (edge (B1, B2, F)),
20 fail
21 ; true
22 ),
23 % findall(edge(I,J,W), edge(I,J,W), Edges),
24 findall(I-J, (edge(I,J,_); edge(J,I,_)), Pairs0),
25 keysort (Pairs0, Pairs1),
26 keyclumped(Pairs1, Pairs2),
27 tell(‘/tmp/all.lp ‘ ),
28 write(‘ Maximize\n obj: ‘ ),
29 (foreach (I1-J1, Pairs1),
30     fromto(‘ ‘,S1,S2,_)
31 do  (I1>J1 -> S1 = S2
32     ; edge (11, J1, F1),
33    format(′~a~w X_~d_~d′, [S1,F1,I1,J1]),
34    S2 = ‘ + ‘
35      )
36 ), nl,
37 write(‘ Subject To\n ‘ ),
38 (foreach (I2-J2s,Pairs2),
39     param(M)
40 do (length(J2s,Len), Len =< M -> true
41     ;   (foreach (J2, J2s),
42        fromto (‘ ‘,S3, ‘ + ‘,_),
43        param(I2)
44   do sort2(I2, J2, I3, J3),
45      format (‘ ~aX_~d_~d ‘, [S3,I3,J3])
46   ),
47   format (‘ <= ~d\n ‘, [M])
48     )
49 ),
50 write(‘ Binary\n ‘ ),
51 (foreach (I4-J4, Pairs1)
52 do  (I4>J4 -> true
53     ;       format (‘ X_~d_~d\n’, [I4,J4])
54     )
55 ),
56 write (‘ End\n ‘ ),
57 told,
58 system(′gurobi_cl ResultFile=/tmp/all.sol /tmp/all.lp′),
59 parse_solution.
63
60  parse_solution :-
61    see(‘/tmp/all.sol ‘ ),
62  retractall(solution(_,_,_)),
63  read_line (_),
64  read_line (_),
65  repeat,
66    read_line (Codes ),
67    (Codes = end_of_file -> true
68    ;   tok_sol_line (I, J, 1, Codes, []),
69        edge (I, J, W),
70        assertz(solution(I,J,W)),
71        fail
72 ),    !,
73  seen,
74  findall(W, solution (_,_,W), Ws),
75  length(Ws, Size),
76  sumlist(Ws, Weight),
77  format (‘ % Gurobi computed a set of ~d edges and weight ~w\n’,
78    [Size, Weight ]),
79  findall(0, (solution(I,J,W), print_tab([I,J,W])), _).
84
80 tok_sol_line (I, J, Z01) --> “X_”,
81   tok_int(0, I), “_”,
82   tok_int (0, J), “ “,
83   tok_int(0, Z01).
89
84 tok_int (Int0, Int) --> [D], {D >= 0 ‘ 0, D =< 0 ‘ 9}, !,
85   {Int1 is 10*Int0 + D - 0 ‘ 0},
86   tok_int (Int1, Int).
87 tok_int (Int, Int) --> [].
94
88 print_tab (L) :-
89   (foreach (X,L) do write(X), write(‘ ‘ )),
90   nl .
98
91 parse_record(Record, B1, B2, F) :-
92   Record = [integer(B1,_), integer(B2,_), float (F,_)].
101
93 bin2chr(Bin, Species, Chr) :-
94   chromosome (Species, Chr, A, B),
95   Bin >= A,
96   Bin =< B, !.
106
97 sort2(X, Y, X, Y) :- X =< Y, !.
98 sort2 (X, Y, Y, X).
~~~

## B Example Data Files

Example Data File 4: An example data file (`.dzn` file) depicting the integral interaction frequencies from the hypothetical whole-genome contact map depicted in Figure 3A utilized
by the **CP** model.

~~~
1 N = 6;
2 map = [|0, 5, 2, 1, 1, 1,
3        |5, 0, 4, 4, 1, 1,
4        |2, 4, 0, 3, 5, 2,
5        |1, 4, 3, 0, 6, 4,
6        |1, 1, 5, 6, 0, 4,
7        |1, 1, 2, 4, 4, 0,
8        |];
~~~

Example Data File 5: An example data file (`testMap.csv`) depicting the interaction frequencies from the hypothetical whole-genome contact map depicted in Figure 3A utilized by the **GM** and **IP** models.

~~~
1 bin_label_1, bin_label_2, interaction_frequency
2  1, 2, 0.5
3  1, 3, 0.2
4  1, 4, 0.1
5  1, 5, 0.1
6  1, 6, 0.1
7  2, 3, 0.4
8  2, 4, 0.4
9  2, 5, 0.1
10 2, 6, 0.1
11 3, 4, 0.3
12 3, 5, 0.5
13 3, 6, 0.2
14 4, 5, 0.6
15 4, 6, 0.4
16 5, 6, 0.4
~~~

## Footnotes

1 http://www.algorithmic-solutions.com/

2 https://developers.google.com/optimization/

3 http://www.minizinc.org/doc-lib/doc-globals-channeling.html

4 http://sicstus.sics.se

5 http://www.gurobi.com/

